# β-glucan induced trained immunity enhances antibody levels in a vaccination model in mice

**DOI:** 10.1101/2024.04.11.588932

**Authors:** Jainu Ajit, Qing Chen, Trevor Ung, Matthew Rosenberger, Jeremiah Kim, Ani Solanki, Jingjing Shen, Aaron P. Esser Kahn

## Abstract

Trained immunity improves disease resistance by strengthening our first line of defense, the innate immune system. Innate immune cells, predominantly macrophages, are epigenetically and metabolically rewired by β-glucan, a fungal cell wall component, to induce trained immunity. These trained macrophages exhibit increased co-stimulatory marker expression and altered cytokine production. Signaling changes from antigen-presenting cells, including macrophages, polarize T-cell responses. Recent work has shown that trained immunity can generally enhance protection against infection, and some work has shown increased protection with specific vaccines. It has been hypothesized that the trained cells themselves potentially modulate adaptive immunity in the context of vaccines. However, the mechanistic link between trained immunity on subsequent vaccinations to enhance antibody levels has not yet been identified. We report that trained immunity induced by a single dose of β-glucan increased antigen presentation in bone-marrow-derived macrophages (BMDMs) and CD4^+^ T cell proliferation *in-vitro*. Mice trained with a single dose of β-glucan a week before vaccination elicited higher antigen-specific antibody levels than untrained mice. Further experiments validate that macrophages mediate this increase. This effect persisted even after vaccinations with 100 times less antigen in trained mice. We report β-glucan training as a novel prophylactic method to enhance the effect of subsequent vaccines.

## Introduction

The immune system consists of innate and adaptive arms, which are known to be linked; however, it has long been thought that only adaptive immunity contains memory.^1,2^ In recent work, innate cells have been revealed to possess nonspecific memory against certain whole-cell vaccines and fungal cell component-β-glucan.^3^ This process, called trained immunity, improves host defense against various bacterial, fungal, and viral infections in mice and humans.^4–6^

Most identified trained immunity-inducing molecules, including bacillus Calmette-Guerin (BCG) vaccine and β-glucan, are also well-known adjuvants.^7,8^ Components of the BCG vaccine activate nucleotide-binding oligomerization domain-2 (NOD 2) receptor, whereas β-glucan binds to the Dectin-1 receptor.^9,10^ Activation of these pattern recognition receptors (PRRs) on innate immune cells induces signaling cascades and affects the adaptive immune response.^11^ In the context of vaccination and mediating adaptive responses, BCG-mediated training of splenic macrophages results in the increased production of pro-inflammatory cytokines like IL-6, TNF-a, and IL-1β and the expression of co-stimulatory markers, such as CD80, CD86, and CD40.^12^ Therefore, it has been hypothesized that trained immunity can improve adaptive immune responses to vaccines and vaccine adjuvants by promoting more robust innate activation.^13,14^ However, there have been limited correlative reports demonstrating the effects of training in protection. These lack a direct testing of how trained antigen-presenting cells induce T-cell activation and antibody production.^15–19^

β-glucan training epigenetically and metabolically rewires many innate immune cell types, including monocytes, macrophages, and neutrophils – ranging in effect from a few days to several months.^20–22^ The difference is due to the differential induction of peripheral or central training.^23^ Short-term training occurs in the periphery, where blood monocytes and tissue macrophages are the critical mediators. The average half-life of these cells is 6-7 days, after which the effects of training subside. Long-term training is mediated by bone marrow progenitors.^21^

Here we investigated if β-glucan training enhances antibody levels in a vaccination model in mice. We observed increased antibody levels in β-glucan trained mice vaccinated with model antigen ovalbumin co-administered with either MPLA (TLR 4 agonist) or Pam3 (TLR 2 agonist). We tested this *in-vitro* and saw that this enhanced effect was driven by trained bone-marrow derived macrophages (BMDMs) capable of robust *in-vitro* CD4^+^ T-cell proliferation. Additionally, we observed higher co-stimulatory marker expression on macrophages in the draining lymph node. We further report that β-glucan induced training enhanced antibody levels with antigen dose sparing up to two orders of magnitude. We also observed higher total IgG in trained mice vaccinated with hemagglutinin, demonstrating the potential of this approach. However, our work in this report only covers a short period of training, and much remains to be learned about longer-term training and its effects on vaccination.

## Results and discussion

### in-vitro β-glucan training in antigen presentation

BCG and adenovirus induce trained immunity resulting in the increased expression of co-stimulatory markers like CD86, CD40, and MHC II.^12,35^ We wanted to confirm that β-glucan training could also induce similar effects on co-stimulatory marker expression to promote antigen presentation. Previous reports showed that β-glucan adjuvanticity increases the expression of co-stimulatory molecules like CD86 and CD40 in dendritic cells via direct activation of Dectin-1, however, no training induced effects have been reported.^24,25^ Therefore, we first tested if β-glucan training improved the expression of co-stimulatory molecules such as CD86 and CD40 on BMDMs in an *in-vitro* training model. In this experiment, 3 million BMDMs were trained with either PBS (untrained) or 100 μg/mL of β-glucan for 24 h in a 12-well plate with a total volume of 2 mL. Then the cells were washed, and the media was replaced. Replacing the media with fresh media ensured no further Dectin-1 stimulation. The cells were then allowed to rest for four days, characteristic of an *in-vitro* training assay. We hypothesized that β-glucan induced training would enhance the expression of co-stimulatory markers like CD86 and CD40 on BMDMs. To test this, cells were analyzed after the four-day resting period by flow cytometry. We observed a 1.3-fold increase in CD86 expression and a 1.8-fold increase in CD40 expression in trained BMDMs (**Fig. 1a and 1b)**. Since training also improves innate cell responses to a secondary adjuvant, a separate set of trained BMDMs was stimulated with Pam3 (TLR 2 agonist) on day 4 and analyzed for co-stimulatory marker expression 24 h later. While CD86 expression increased 1.8 fold, CD40 expression enhanced 1.1 fold after Pam3 stimulation (**Fig. 1a and 1b)**. These results demonstrated that β-glucan induced training improved macrophage antigen presentation.

**Fig 1:**
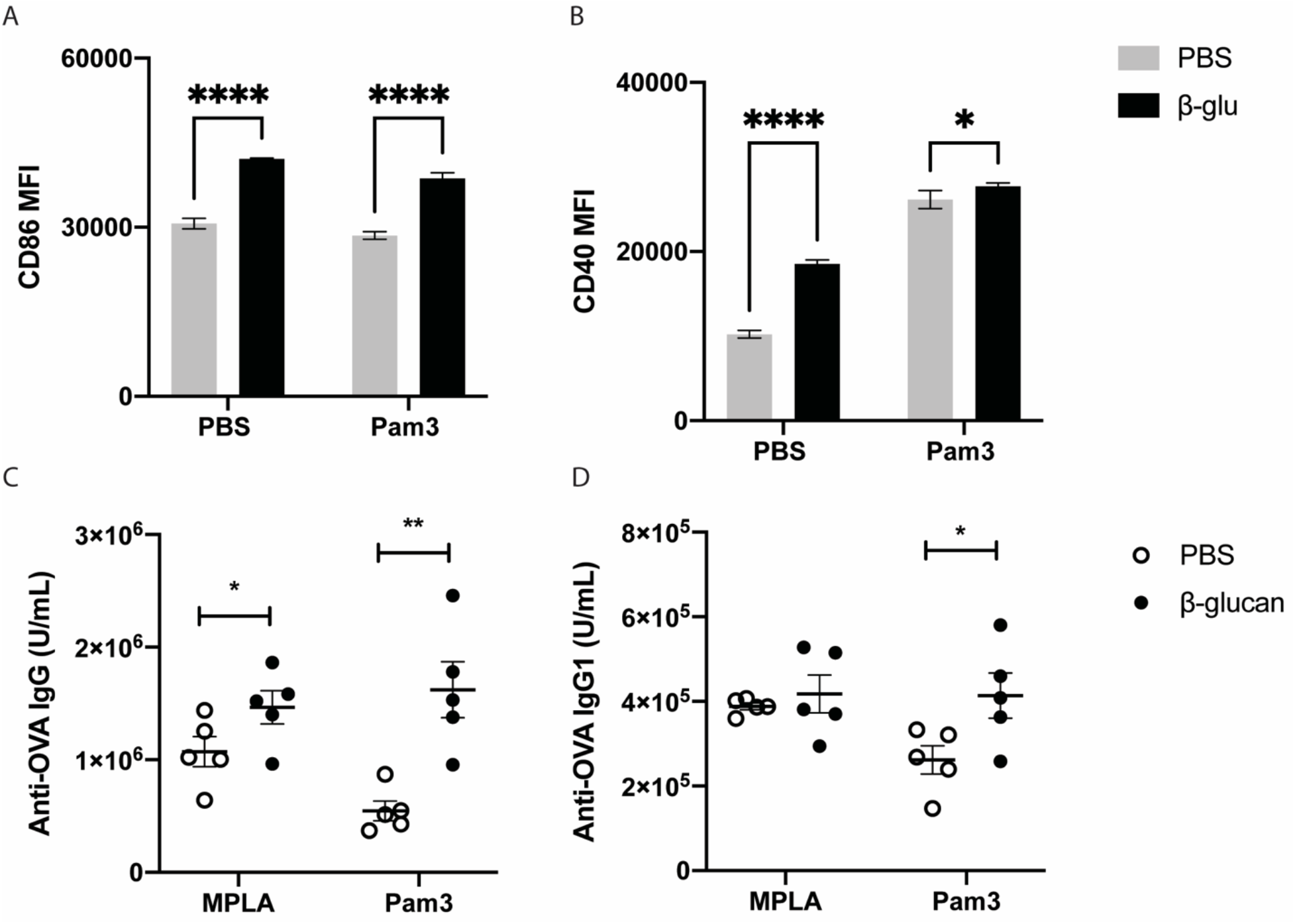
*in-vitro* β-glucan training-induced enhanced antigen presentation in BMDMs and in-vivo vaccination with model antigen OVA. BMDMs were trained with PBS (grey bar) or 100 μg/mL of β-glucan (black) for 24 h. Cells were then washed and rested for four days. Cells were released from the plate on day 5 and analyzed for expression of A) CD86 or B) CD40 by flow cytometry. Mice were trained i.p. with PBS (grey) or β-glucan (black). 1 week later, mice were vaccinated i.m containing OVA (100 μg) with either MPLA or Pam3 (10 μg). Mice were boosted 2 weeks later, and serum cytokines were analyzed for antibody levels C) IgG and D) IgG1 2 weeks post boost. Statistics were calculated using student’s T test ; n=5; *P < 0.05, **P < 0.01, and ***P < 0.001. n.s., not significant.

### in-vivo β-glucan training in improving the effect of vaccination

Enhanced antigen presentation stimulates T-cell proliferation and increases the production of antigen-specific antibodies. We next performed a vaccination experiment to determine if the enhanced expression of co-stimulatory markers on BMDMs observed *in-vitro* would be observed for mice experiencing training *in-vivo*. We tested if a training regimen, followed by a model vaccination, would result in improved antibody levels *in-vivo*. In this experiment, mice were trained intraperitoneally with PBS (untrained) or 1 mg of β-glucan. Mice were allowed to rest for one week to ensure that all training material was dispersed and did not contribute to adjuvanticity. One week post-training, mice were vaccinated subcutaneously with model antigen ovalbumin (OVA) co-administered with a TLR agonist. A subcutaneous route of administration was chosen for vaccinating the animals to minimize any potential adjuvant effects of persistent β-glucan. We performed a preliminary screen to determine the adjuvant that would work best with the training model considering possible synergistic interactions. In this screen, we utilized Pam3CSK4 (TLR1/2 agonist), MPLA (TLR4 agonist), R848 (TLR7/8 agonist), and CpG (TLR 9 agonist). OVA was administered at 100 μg per mouse, and adjuvant concentrations were determined based on literature precedent.^26^ (See materials and methods for details) Vaccinated mice received an identical boost two weeks later. Antibody levels were analyzed two weeks after the boost via retro-orbital bleed. We observed significantly higher anti-OVA IgG and IgG_1_ levels in β-glucan trained mice that were subsequently vaccinated with OVA and MPLA, and Pam3CSK4. (**Fig. 1c, 1d**) No difference was observed with the other adjuvants tested (**Fig. SI 1)**. This observation agreed with prior literature, citing higher trained immunity effects due to increased synergies between Dectin-1 and TLR2 and TLR4 mediated immune activation.^27^ Taken together, these results suggest that β-glucan training enhances antibody levels when combined with the appropriate agonist/antigen combinations.

### Mechanism of action

β-glucan also binds to Dectin-1 on B-cells to upregulate pro-inflammatory cytokines such as IL-6 and TNF-α.^28^ To ensure that β-glucan mediated enhancement in antibody production is mediated solely via innate immune cells and subsequent antigen presentation, we performed the same vaccination experiment in MHC II knockout mice. We did not observe measurable levels of antibodies in MHCII knockout mice, indicating that potential β-glucan activation or proliferation of B-cells did not contribute to an increase in antibodies. (**Fig. SI 2**)

Trained immunity can be elicited at two levels: peripheral training by tissue-resident innate cells (short-term) and central training by bone-marrow progenitors (long-term).^29^ We sought to better understand which of these two types of training contribute to the mechanism of action of β-glucan induced enhancement in antibody responses. We compared different routes of administration of training and vaccination while maintaining the timing of experiments (**Fig. 2a**). Intra-peritoneal training leads to central training by promoting myelopoiesis in progenitor cells from the bone marrow and spleen.^30–33^ We hypothesized that central training would contribute to improved antibody levels irrespective of the route of vaccination. To test this, we fixed the route of training (intra-peritoneal) and changed the route of vaccination from subcutaneous to intramuscular. If the effects of β-glucan training occurred centrally, the route of vaccination would have minimal effects on the antibody titer. Indeed, we observed that intramuscular vaccination also enhanced antibody levels in β−glucan trained mice. (**Fig. 2b**) To examine if inducing peripheral training in the draining lymph nodes enhanced antibody levels, we localized training and vaccine regimens by performing both subcutaneously. We hypothesized that in this experiment, peripheral trained antigen-presenting cells would contribute to enhanced T-cell activation and subsequent improvement in antibody levels. However, we observed no significant change in antibody levels when training and vaccination were given at the same site (**Fig 2b**). From these results, we reasoned that the well-established systemic effects following intraperitoneal administration of β-glucan were key in determining antibody levels.

**Fig 2:**
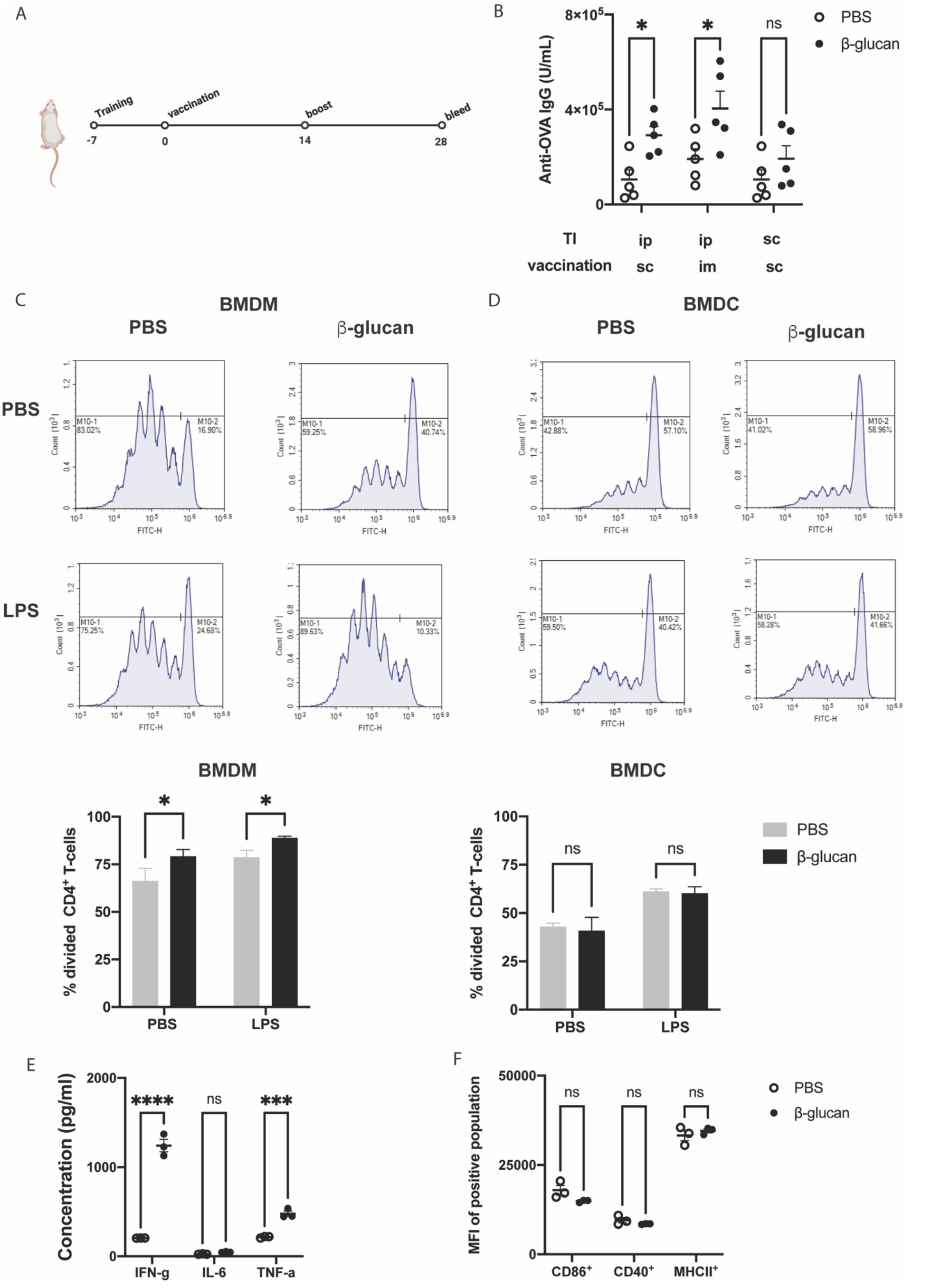
Mechanism of β−glucan training induced enhancement in antibody levels. a) *in-vivo* vaccination schedule with model antigen ovalbumin. b) Mice were trained with PBS (grey) or β-glucan (black). 1 week later, mice were vaccinated containing OVA with Pam3. Mice were boosted 2 weeks later, and serum cytokines were analyzed for anti-OVA IgG. c) BMDMs and d) BMDCs were harvested from mice trained with PBS (grey bar) or 100 μg/mL of β-glucan (black). They were co-cultured with FITC-labeled CD4^+^ T cells in media alone or with LPS for 5 days, after which T-cell proliferation was accessed. e) Supernatant from the co-culture assay with BMDMs and CD4+T cells were analyzed. f) BMDMs were isolated from PBS (grey) or β-glucan (black) trained mice after 5 days and were analyzed for the expression of co-stimulatory molecules. Statistics were calculated using student’s T test or 2-way ANOVA ; n=5; *P < 0.05, **P < 0.01, and ***P < 0.001. n.s., not significant.

After determining that central training mediated enhanced response to vaccination following β-glucan induced training, we sought to determine which cell types were responsible. β-glucan central training occurs mainly in the bone marrow and spleen and primarily in macrophages.^20,34^ To identify cells, we trained mice with the standard dose of 1 mg β-glucan intraperitoneally. After seven days, we harvested splenic macrophages and bone marrow. Bone-marrow cells were differentiated into macrophages and dendritic cells separately using MCSF and GMCSF, respectively, in culture media. To test which cells enhanced antigen presentation, we performed an *in-vitro* co-culture assay with FITC labeled CD4^+^ T cells from OT-II mice for five days with or without agonists - LPS or Pam3. At the end of five days, we evaluated T-cell proliferation and cytokine production in the supernatant (Gating strategy in **Fig. SI 3**). We observed that only bone marrow derived macrophages (BMDMs) from β-glucan trained mice induced significantly higher CD4^+^ T cell proliferation *in-vitro* (**Fig. 2c**). Bone Marrow Derived Dendritic Cells (BMDCs) and splenic macrophages from β-glucan trained mice did not induce any significant improvement in T-cell proliferation (**Fig. 2d** and **Fig. SI 4**). This provides another piece of evidence that cells in the bone marrow are partially responsible for the increased response, and their activity is enhanced by LPS though potentially not entirely due to it. To determine the mechanism by which T-cells were expanding, cytokines from the co-culture supernatant was analyzed. Cytokine analysis revealed that β-glucan trained BMDMs increased IFN-γ and TNF-α expression when co-cultured with T-cells in the presence of LPS, indicating a TH1 response (**Fig 2e**). To determine whether this response might correlate with co-stimulatory marker expression, BMDMs from PBS and β-glucan trained mice were analyzed by flow cytometry. However, we observed no significant differences in the expression level of co-stimulatory markers between untrained and β-glucan BMDMs (**Fig 2f**). Taken together, these results suggested that β-glucan induced trained immunity reprogrammed bone marrow macrophages to elicit higher CD4^+^ T cell proliferation.

Based on this data, we hypothesized that T-cell responses might be enhanced in β-glucan trained mice vaccinated with ovalbumin. To test this, we repeated the previous vaccination schedule (**Fig. 3a**), inducing training with β-glucan a week before vaccinating intramuscularly with ovalbumin. We used Pam3 as the agonist of choice due to higher antibody levels observed in the previous experiment. (**Fig. 1d**) Two weeks after the boost, we harvested the draining inguinal lymph node and spleen to analyze antigen-specific CD4^+^ T-cell responses. We observed a non-significant increase in OVA-specific MHCII tetramers in splenocytes isolated from β-glucan trained mice (**Fig. 3a**). Intracellular staining of CD4^+^ T-cells from lymphocytes and splenocytes revealed no significant differences between untrained and trained mice in IL-4 levels (**Fig. SI 5**). We also analyzed the antigen recall activity of isolated splenocytes to OVA at a concentration of 1 μg/mL. We observed higher levels of IFN-γ (**Fig. 3b**) and IL-2 (**Fig. SI 6**) from splenocytes isolated from β-glucan trained mice compared to untrained controls. These results demonstrated that β-glucan training induced an antigen-specific TH1 response in draining lymph nodes and the spleen. However, it appears that this response may not lead to a significant increase in the number of antigen-specific CD4^+^ T-cells, but only in their responsiveness to antigen.

**Fig 3:**
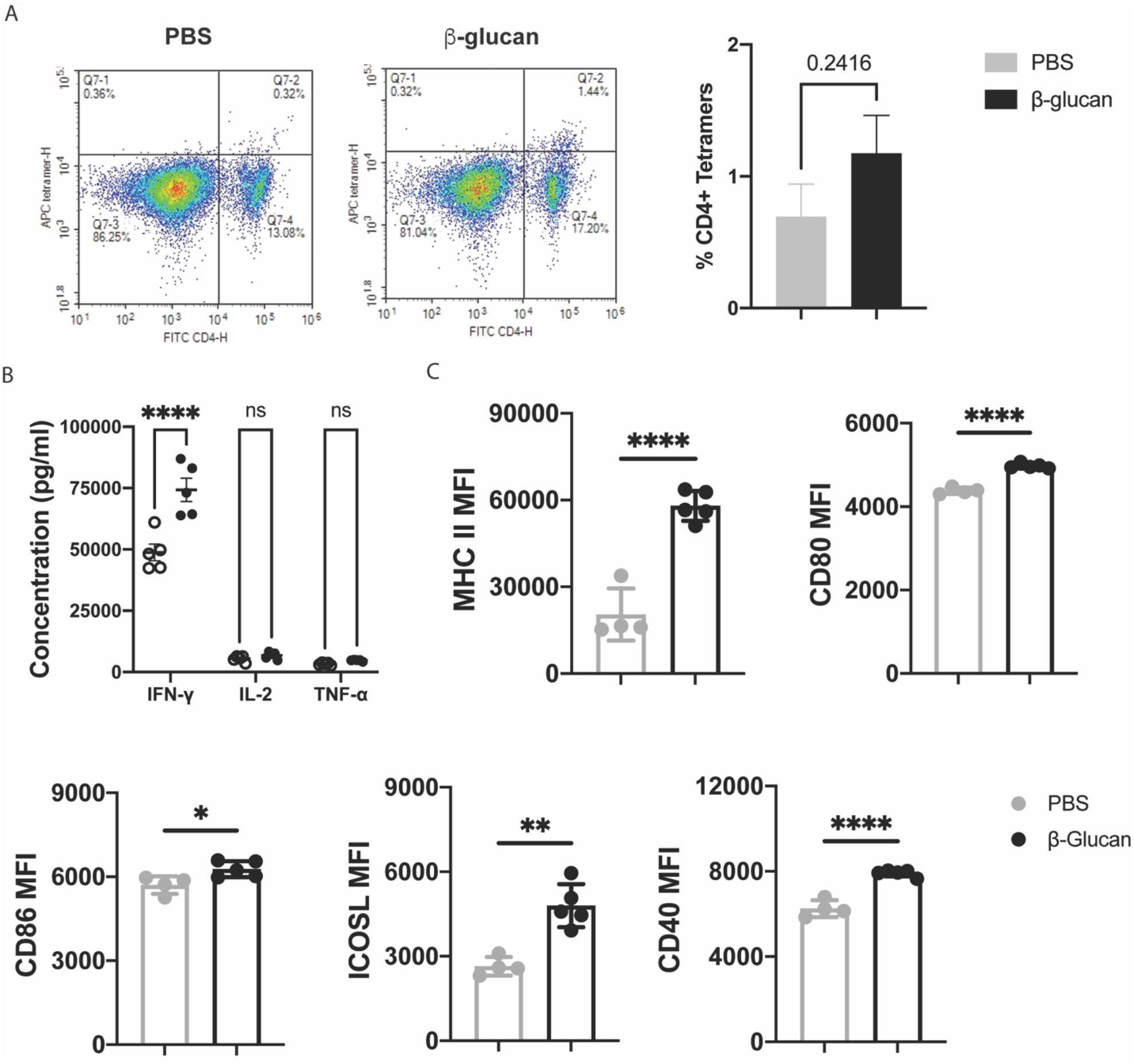
Determining the cellular players of β-glucan training induced enhancement in antibody levels. a) OVA-specific CD4+ T cell analysis on day 28 using MHC-II tetramer assay from splenocytes isolated from mice trained with PBS (grey) or β-glucan (black). b) Antigen recall assay from splenocytes isolated on day 28 from PBS (white) or β-glucan (black) trained mice. c) Anti-OVA titer on day 49 from PBS (white) or β-glucan (black) trained mice. d) day 42 draining lymph node analysis for the expression of co-stimulatory markers from PBS (green) or β-glucan trained mice (red). Statistics were calculated using student’s T test or 2-way ANOVA ; n=5; *P < 0.05, **P < 0.01, and ***P < 0.001. n.s., not significant.

Next, we tested to see if β-glucan training enhanced antibody levels over longer durations of time-42 days after the first vaccination. We performed the same experiment as described before (**Fig. 2a**), measured antibody titers, and analyzed cell populations in the draining lymph node on day 42 (**Fig. SI 7**). We observed a continued 1.5-fold increase in antibody levels even after 42 days (**Fig. SI 8**). Since we observed increased T-cell proliferation by β-glucan-trained BMDMs, we hypothesized that macrophages in the draining lymph node would exhibit higher expression of co-stimulatory markers. Analysis of macrophages in the draining lymph node revealed upregulation in CD40, CD80, CD86, ICOSL, and MHC II expression (**Fig. 3c**) This further supported our observation that macrophages play a key role in enhanced antibody responses in β-glucan trained mice.

### Application

An important aspect of vaccination is dose-sparing, the reduction of antigen use. As training resulted in cells capable of greater antigen presentation, we tested if β-glucan induced training lowered the amount of antigen needed to produce robust antigen-specific antibody responses. We reduced the dose of ovalbumin from 100 μg down to 1 μg per mouse and followed the same vaccination schedule with Pam3 as the agonist of choice (**Fig. 2a**). We observed a consistent and statistically significant enhancement in antibody levels from β-glucan trained mice. Most notably, we observed a 1.6-fold increase in antibody levels in β-glucan trained mice vaccinated with just 1 μg OVA compared to untrained controls (**Fig. 4a**)

**Fig 4:**
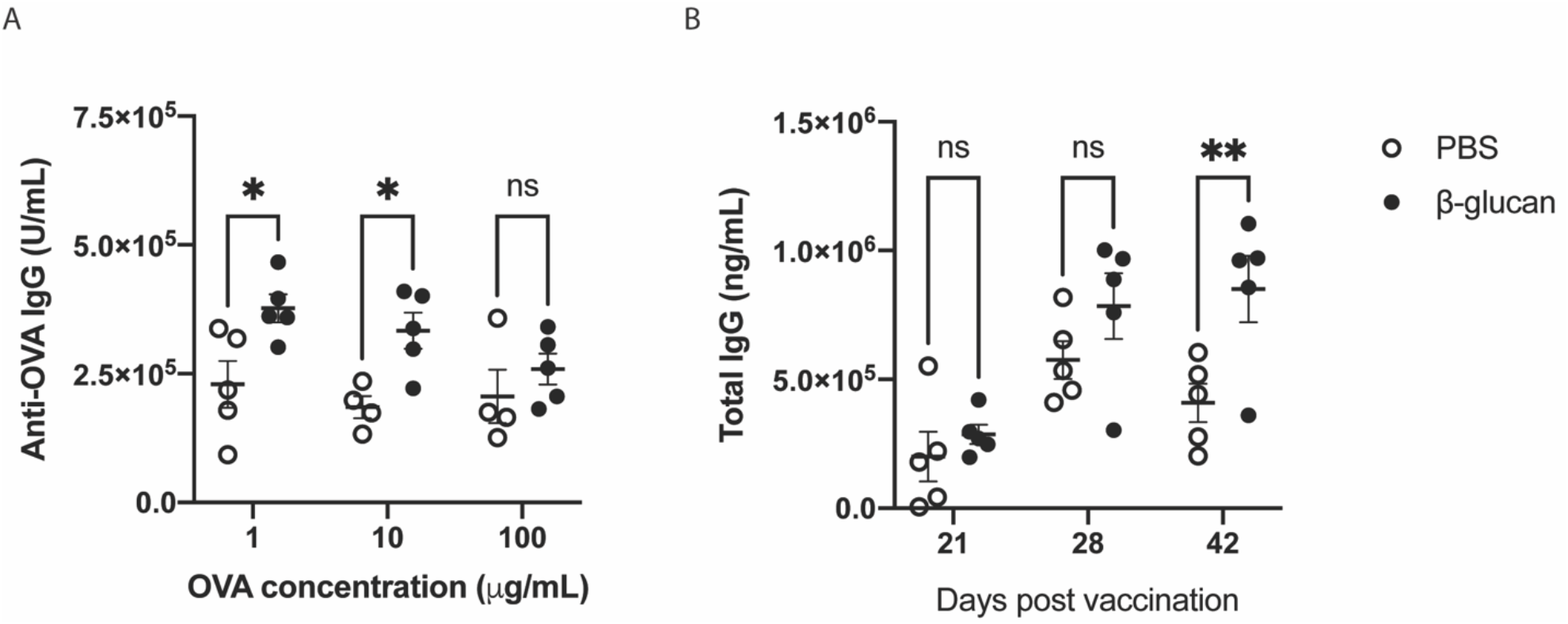
Applications of β-glucan training in prolonged and broad-range vaccinations. a) Anti-OVA titer on day 28 from PBS (white) or β-glucan (black) trained mice vaccinated intramuscularly with varying doses of ovalbumin with 10 μg of Pam3. b) Total IgG titer from PBS (white) or β-glucan (black) trained mice vaccinated intramuscularly with HA protein and Pam3 at 21, 28, and 42 days post-vaccination. Statistics were calculated 2-way ANOVA ; n=5; *P < 0.05, **P < 0.01, and ***P < 0.001. n.s., not significant.

We next wanted to test if β-glucan training enhances antibody levels to other antigens. We selected influenza as a proof-of-concept vaccine due to its relevance in current infections that require seasonal shots. We hypothesized that trained immunity could enhance antibody titers and thereby decrease or eliminate the need for seasonal vaccinations. Mice were vaccinated with Influenza A H1N1 (A/California/04/2009) hemagglutinin (HA) protein adjuvanted with Pam3. Mice were trained according to our existing protocol (**Fig. 2a**), and total IgG was measured from the serum of mice on days 21, 28, and 42 post-vaccination. We continued to observe higher total IgG from β-glucan trained mice compared to untrained controls. Additionally, we observed a 2-fold increase in total IgG that persisted until 42 days post-vaccination (**Fig. 4b**). This result provides preliminary proof of concept that trained immunity can be harnessed to improve antibody levels for different antigens at use in subunit vaccines.

## Discussion

In this work, we examined the role of trained immunity in promoting an increased adaptive response against select antigens and model vaccinations. Innate immune cells shape adaptive responses through cytokines and co-stimulatory molecule expression. Trained immunity induces epigenetic and metabolic reprogramming of macrophages, facilitating higher inflammatory cytokines and cell surface markers. Previous work on BCG and adenovirus demonstrated the increased expression of co-stimulatory markers-MHCII, CD40, and CD80 in macrophages.^12,35^ BCG vaccine induces heterologous TH1/TH17 activation in humans. These key signals play essential roles in polarizing T-cell responses. However, how these effects translate to increased protection in combination with other vaccinations is not fully understood. Moreover, the therapeutic potential of employing trained immunity to act as a pre-vaccination “coach” to enhance innate responses has had limited exploration. Here, we report the enhanced antibody responses generated from training mice with β-glucan a week before vaccination.

β-glucan training improved antigen presentation in BMDMs *in-vitro* and enhanced OVA-specific antibody levels. To identify the cells responsible for this observed enhancement, we isolated BMDMs from mice trained with β-glucan, which induced higher CD4^+^ T cell proliferation than untrained controls. In addition, supernatant from the co-culture assay showed elevated IFN-γ and TNF-α levels indicative of a TH1 response. Finally, β-glucan training improved dose sparing and generated robust antigen-specific antibodies with very low antigen dosage (100-fold less). Moreover, we show that training can be used to enhance the effects of vaccinations with other antigens, such as hemagglutinin. The results indicated that the enhancement from training is mediated by BMDMs and not DCs or splenocytes. This intriguing mechanism opens the potential for further investigation of improved training of BMDMs or seeking methods to train other cell populations. These results demonstrate that trained immunity can be used as an enhancing agent to safely improve the effects of a potentially broad range of vaccines with increased antibody responses. In prior work, training phenomena have resulted in increased protective effects of other vaccination. These results provide a potential mechanistic pathway by which macrophages assist in promoting improved vaccine responses. This intriguing hypothesis will require a great deal more exploration.

These results also indicate the potential to design novel materials that target training in addition to conventional signaling pathways. β-glucan conjugated materials improve antigen presentation and CD4 T-cell proliferation and merits further exploration.^38,39^ Novel therapies might be designed to specifically train macrophages prior to vaccinations.

## Materials and Methods

All chemicals and reagents unless noted otherwise were purchased from Sigma. ELISA kits and all fluorescently tagged antibodies were purchased from BioLegend. MHCII Tetramers (OVA323-339 –PE conjugate was purchased from MBL International (Woburn, MA). ODN 1826 (CpG), Pam3, MPLA and alum were purchased from Invivogen. All cell culture reagents were obtained from Thermo Fisher Scientific. Cells were maintained at 37 °C and 5% CO2. C57Bl/6J mice were obtained from Jackson Laboratories and acclimatized for 1 week prior to experimentation. All animal experiments were conducted with approval from the University of Chicago Institutional Animal Care and Use Committee (approval number 72517). All statistical analyses were performed using GraphPad Prism.

### *In-vitro* training assay

BMDMs were plated at a density of 3*10^6 cells/well in flat bottom 12-well plates (Corning) at a final volume of 2 mL) and rested for a few hours to adhere at 37 ºC and 5% CO_2_. After the cells were adherent, training material was added at the desired concentration and incubated for 24 h. Then cells were washed and rested for 3 days. On day 4, BMDMs were released and stained with blocking antibody (CD16/32) for 10 min. The cells were then stained with anti-mouse FITC-CD86, PE-CD40, or APC-MHC II and analyzed using flow cytometry. A separate set of trained cells were stimulated with 10 ng/mL of Pam3 overnight and analyzed by flow cytometry using ACEA Novocyte after 18 h.

### Vaccination experiment and antibody quantification

Mice were lightly anesthetized with isoflurane and injected with 1 mg free β-glucan or sterile PBS (control) in a total volume of 150 uL either i.p or s.c near the spleen, as indicated. After 7 days, mice received vaccination (i.m or s.c) containing OVA (100 μg) with an adjuvant of choice at the following concentrations: MPLA and Pam_3_CSK_4_ at 10 μg/mouse, CpG and R848 at 50 mg/mouse or Alum (1:1 with antigen OVA). For hemagglutinin vaccination, A/Michigan/45/2015 X-275 (H1N1) (Sino Biological) was used at a concentration of 5 μg per mouse. Mice received the same formulation as a boost two weeks later. Blood was collected at time points indicated in 0.2 ml of heparin-coated collection tubes (VWR Scientific) for plasma or uncoated tubes for serum. Plasma was isolated via centrifugation (2000*g*, 5 min). Serum was isolated by allowing blood to clot for 15 to 30 min at RT and centrifuging (2000*g* for 10 min) at 4°C. Serum was analyzed using a quantitative anti-OVA total Ig’s ELISA kit (Alpha Diagnostic International) according to the specified protocol. Total IgG was analyzed using total mouse IgG uncoated ELISA (Invitrogen). Plates were analyzed using a Multiskan FC plate reader (Thermo Fisher Scientific), and absorbance was measured at 450 nm and 630 nm. Data were analyzed using GraphPad Prism.

### Cell harvest and culture

Monocytes were harvested from the femurs of 6 week old C57BL/6 mice (Jackson Laboratory). Monocytes were differentiated into macrophages using supplemented culture medium: RPMI 1640 (Life Technologies), 10% heat inactivated fetal bovine serum (HIFBS) 2 × 10^−3^ m L-glutamine (Life Technologies), antibiotic antimycotic (1×) (Life Technologies), and 10% MCSF (mycoplasma free L929 supernatant) for 5 d at 37 °C and 5% CO2. The cells were then released with 5 × 10^−3^ m EDTA in PBS, counted and plated at desired densities. Monocytes were differentiated into dendritic cells using RPMI 1640 (Life Technologies), 10% HIFBS (Sigma-Aldrich), GM-CSF (20 ng/ml; BioLegend), 2 mM l-glutamine (Life Technologies), 1% antibiotic antimycotic (Life Technologies), and 50 μM β-mercaptoethanol (Sigma-Aldrich). Cells were used after 6 days of culture. Spleens were isolated from naive or trained mice. Spleens were homogenized, and cells were filtered through a 70 μm strainer. Red blood cells were lysed by incubating with ACK Lysing Buffer for 5 min at 25 °C. Cells were plated in petri dishes for 24 h and washed to remove non-adherent cells. The adherent cells (splenic macrophages) were used for analysis.

### Antigen recall assay

Mice were vaccinated and boosted on day 14 as described above. On day 28, spleens were harvested and placed in PBS on ice. 1 million cells were added to a flat-bottom 96-well plate precoated with anti-CD3 (3 μg/ml). Antigen or epitope (20 μg/ml) and anti-CD28 (5 μg/ml) were added, and cells were incubated for 48 hours. After 48 hours, cells were pelleted at 400g for 10 min, and cell supernatant was removed and analyzed using LEGENDplex Mouse Th Cytokine Panel according to the manufacturer’s protocol.

### Tetramer Assay

Mice were vaccinated and boosted on day 14, as described above. On day 28, spleens were harvested and placed in PBS on ice. 2 million cells were added to a flat-bottom 96-well plate was incubated with 10 μg/mL OVA 323–339 peptide (major MHCII epitope) for 48 h. The cell culture was supplemented with 0.5 ng/mL IL-2 after 2 days. Cells were analyzed four days following IL-2 stimulation by flow cytometry for CD3+ CD4+ tetramers (Fig SI 5).

### T-cell proliferation assay

T cells were extracted from OT-II mouse spleens and magnetically sorted using a T-cell extraction kit from Stem Cell Technologies (catalog number 19851). Cells were incubated in culture media (RPMI+ 10% HIFBS + Pen/Strep + 1 mM HEPES + 50 μM beta mercaptoethanol) then labelled with CFSE (10 μM) for 20 min. The cells were then washed and incubated at a concentration of 1.5 million cells with 500 k of the selected cell type (e.g. BMDMs, BMDCs) for 5 days. Cells were spun down at 300 g for 5 min and supernatant was collected for analysis by mouse inflammatory cytokine bead array (CBA, BD Biosciences). Cells were resuspended and stained for CD4 and analyzed for proliferation based on CFSE decrease (Fig SI 3).

### Statistical Analysis

All values are expressed as mean ± SEM. The sample size is as indicated in figure captions in all *in-vivo* and *in-vitro* experiments. Student’s T-test was applied for comparing two groups, and one- or two-way analysis of variance (ANOVA) followed by Tukey’s or Dunnett’s multiple comparisons for comparison of multiple groups (indicated in the figure captions) using the GraphPad Prism 9 software. *P* values less than 0.05 were considered statistically significant. Significance \**P* < 0.05, \*\**P* < 0.01, \*\*\**P* < 0.001, \*\*\*\**P* < 0.0001, n.s., not significant.

## Acknowledgements

The authors thank Dr. Emiliano Gomez Medellin for helpful discussions. The authors thank University of Chicago veterinary technicians for exceptional animal care. This work was supported by a grant from DTRA (HDTRA11810052).

## Conflict of Interest

The authors declare no conflict of interest.

### Author contributions

J.A. and A.P.E-K. designed the experiments. J.A. and Q.C. performed the in vitro experiments. J.A., Q.C., T.U., M.R., J.K., A.S. and J.S., performed the in vivo experiments. J.A. and A.P.E.K wrote the manuscript. All authors reviewed and approved the manuscript prior to submission.

## Notes

### Competing Interest Statement

The authors have declared no competing interest.

