## Supplementary material for "β-glucan induced trained immunity enhances antibody levels in a vaccination model in mice": SI

**Fig.SI 1 day 28 Anti-OVA IgGAM and IgG after vaccination**

Mice were trained with PBS (grey) or β-glucan (black) intraperitoneally. 1 week later, mice were vaccinated containing OVA with either MPLA or Pam3. Mice were boosted 2 weeks later and serum cytokines analyzed for antibody levels a) IgGAM and b) IgG 2 weeks post boost. n=5; statistics were calculated using student's T test. **P* < 0.05, ***P* < 0.01, and ****P* < 0.001. n.s., not significant.

**a)**

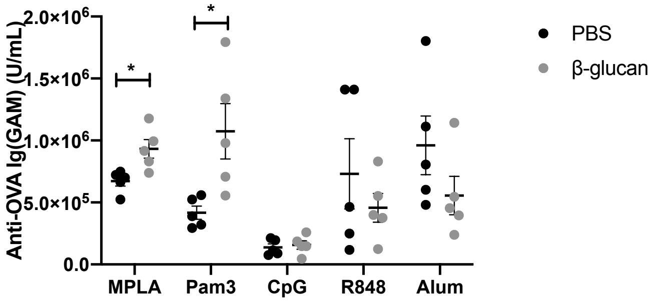


**b)**

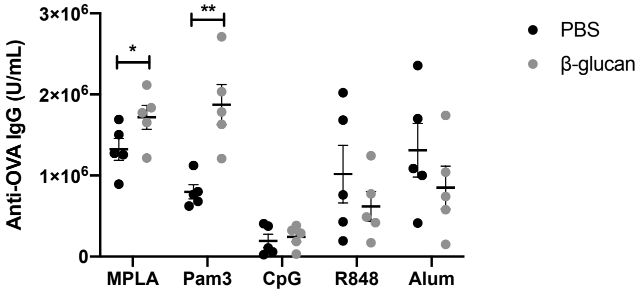


**Fig SI 2: Dependance of MHC II on β-glucan mediated enhancement in immune responses**

Regular C57b6 mice or MHC II $^{-/-}$ mice were trained with PBS (white) or β-glucan (black) intraperitoneally. 1 week later, mice were vaccinated containing OVA with either MPLA or Pam3. Mice were boosted 2 weeks later and serum cytokines analyzed for antibody levels IgGAM 2 weeks post boost. n=5; statistics were calculated using student's T test. **P* < 0.05, ***P* < 0.01, and ****P* < 0.001. n.s., not significant.

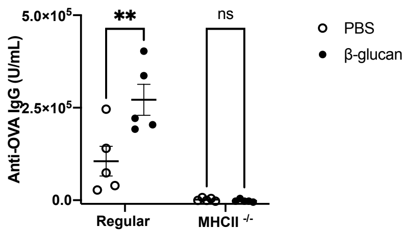

**Fig SI 3: Gating strategy for analysis of CD4$^+$ T cell proliferation**

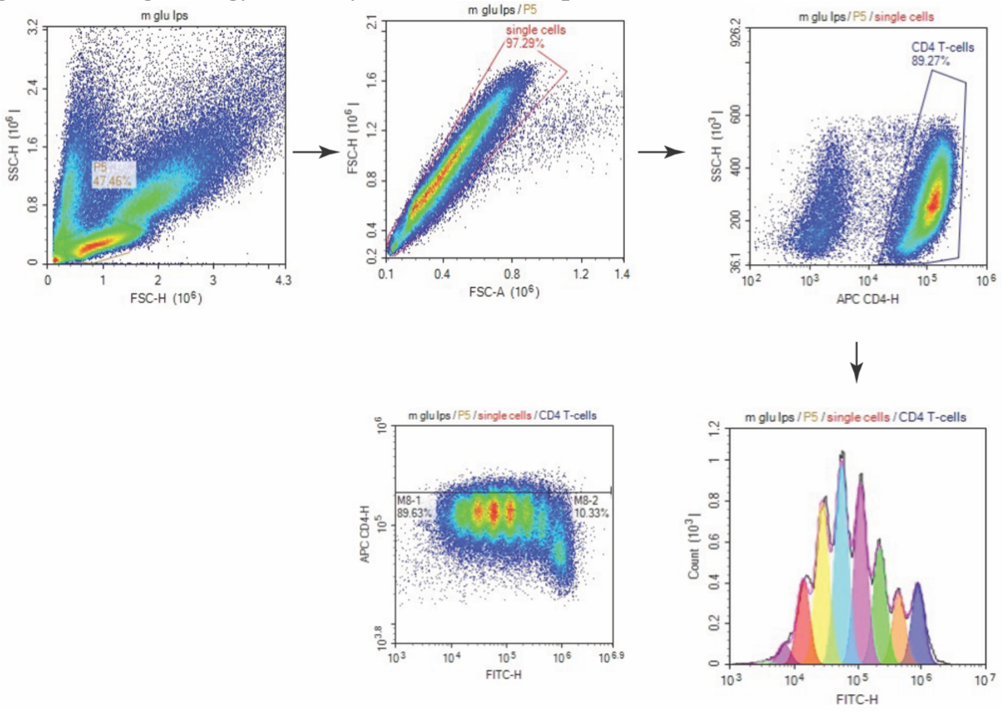


**Fig SI 4: Splenic macrophages does not increase CD4$^+$ T cell proliferation *in-vitro* with LPS**

Splenic macrophages were harvested from mice trained with PBS (grey bar) or 100 µg/mL of β-glucan (black). They were co-cultured with FITC-labeled CD4$^+$ T cells in media alone or with LPS for 5 days, after which T-cell proliferation was accessed. n=5; statistics were calculated using student's T test. *$P < 0.05$, **$P < 0.01$, and ***$P < 0.001$. n.s., not significant.

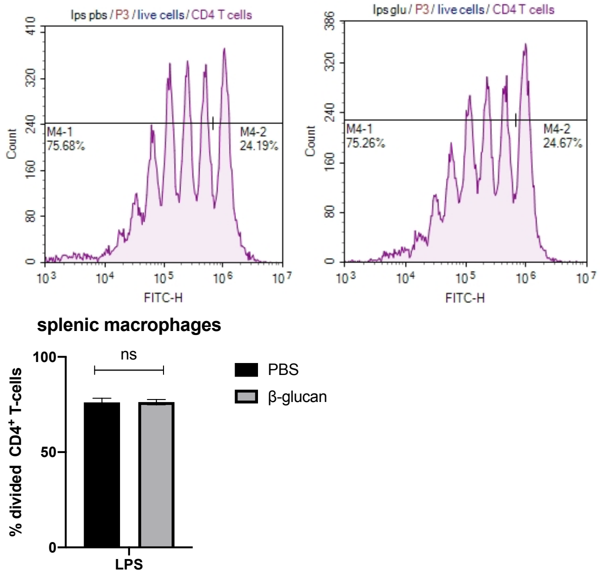


**Fig SI 5: Gating strategy and results for intracellular cytokine staining for CD4$^+$ IL4$^+$ T cells from day 28 splenocytes and lymphocytes**

Intracellular cytokine staining with lymphocytes and splenocytes on day 28 isolated from mice trained with PBS (white) or β-glucan (black). Statistics were calculated using student's T test; n=5; *P < 0.05, **P < 0.01, and ***P < 0.001. n.s., not significant.

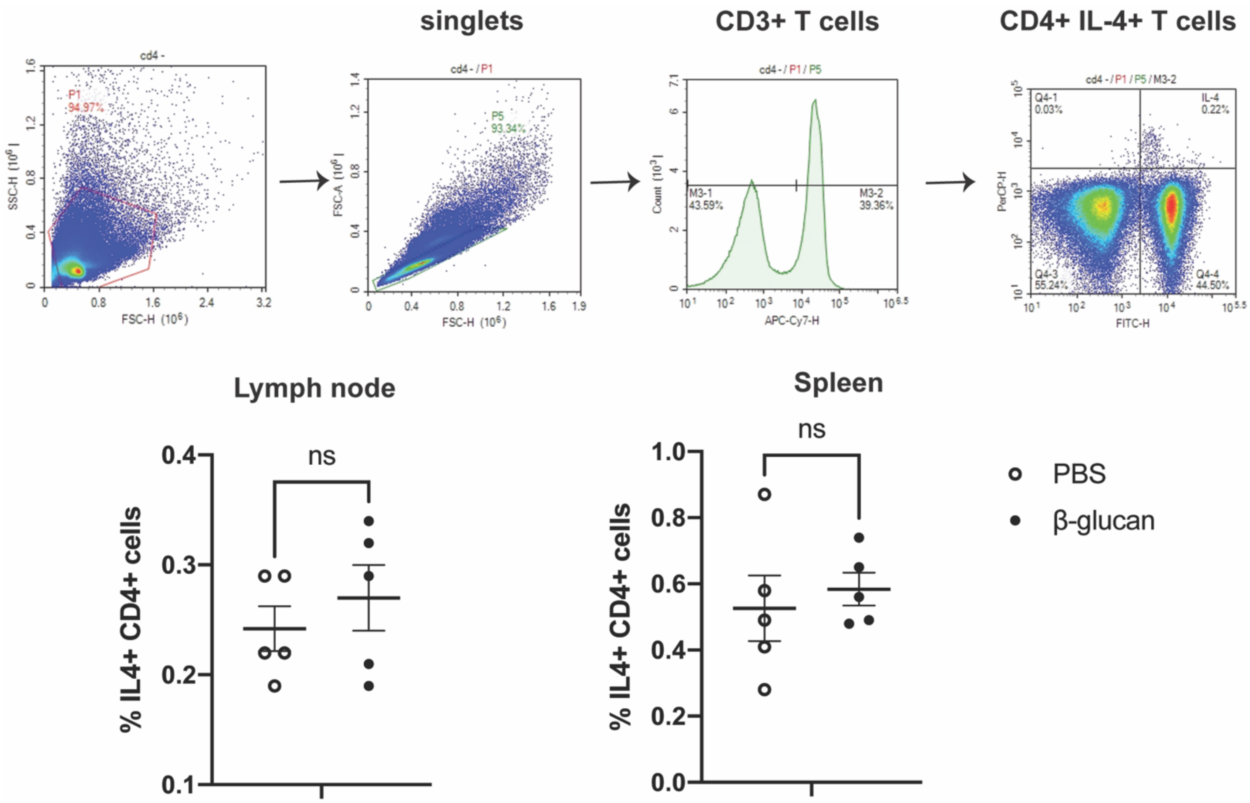


### Fig. SI 6: T-cell recall assay with splenocytes stimulated without anti-CD28 or anti-CD3

Mice were trained with PBS (white) or β-glucan (black) and vaccinated with ova. On day 28, splenocytes were isolated and cultured in 96 well plates with 1 mg/mL OVA for 48 h. Statistics were calculated using student's T test; n=5; *P < 0.05, **P < 0.01, and ***P < 0.001. n.s., not significant.

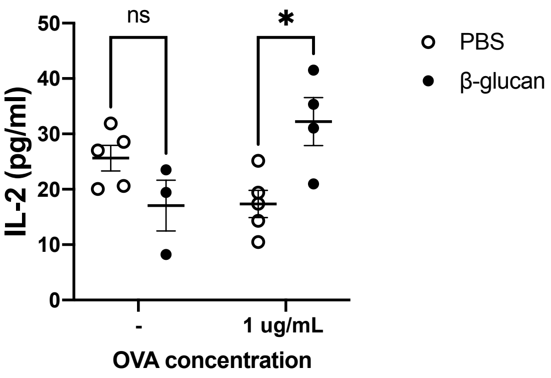

### Fig. SI 7: Day 42 gating strategy and analysis of germinal center B-cells and macrophages in draining lymph node

Mice were trained with PBS (white) or β-glucan (black) and vaccinated with ova. On day 42, draining inguinal lymph nodes were isolated and analyzed for costimulatory markers. Percentage and number of macrophages were evaluated. n=5

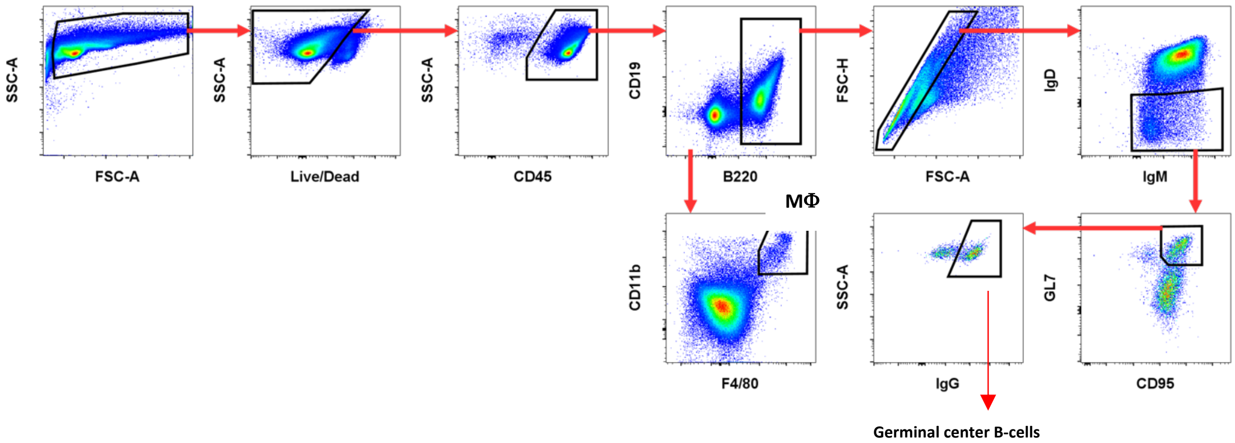


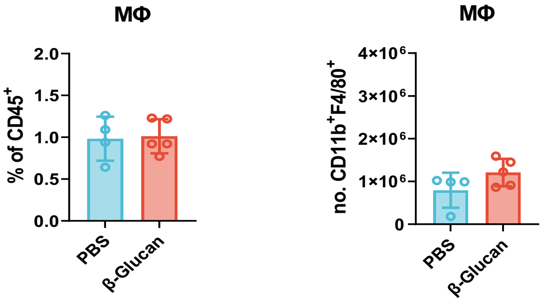


### Fig. SI 8: Day 42 anti- OVA antibody levels

Mice were trained with PBS (white) or β-glucan (black) and vaccinated with ova. On day 42, serum was analyzed for anti-ovalbumin antibodies.

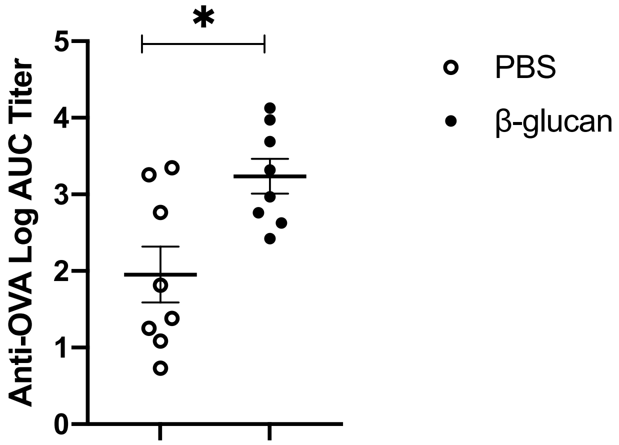
